# XBB.1.5 Spike Protein COVID-19 Vaccine Induces Broadly Neutralizing and Cellular Immune Responses Against EG.5.1 and Emerging XBB Variants

**DOI:** 10.1101/2023.08.30.554497

**Authors:** Nita Patel, Jessica F. Trost, Mimi Guebre-Xabier, Haixia Zhou, Jim Norton, Desheng Jiang, Zhaohui Cai, Mingzhu Zhu, Anthony M. Marchese, Ann M. Greene, Raburn M. Mallory, Raj Kalkeri, Filip Dubovsky, Gale Smith

## Abstract

Monovalent SARS-CoV-2 Prototype (Wuhan-Hu-1) and bivalent (Prototype + BA.4/5) COVID-19 vaccines have demonstrated a waning of vaccine-mediated immunity highlighted by lower neutralizing antibody responses against SARS-CoV-2 Omicron XBB sub-variants. The reduction of humoral immunity due to the rapid evolution of SARS-CoV-2 has signaled the need for an update to vaccine composition. A strain change for all authorized/approved vaccines to a monovalent composition with Omicron subvariant XBB.1.5 has been supported by the WHO, EMA, and FDA. Here, we demonstrate that immunization with a monovalent recombinant spike protein COVID-19 vaccine (Novavax, Inc.) based on the subvariant XBB.1.5 induces cross-neutralizing antibodies against XBB.1.5, XBB.1.16, XBB.2.3, EG.5.1, and XBB.1.16.6 subvariants, promotes higher pseudovirus neutralizing antibody titers than bivalent (Prototype + XBB.1.5) vaccine, induces SARS-CoV-2 spike-specific Th1-biased CD4+ T-cell responses against XBB subvariants, and robustly boosts antibody responses in mice and nonhuman primates primed with a variety of monovalent and bivalent vaccines. Together, these data support updating the Novavax vaccine to a monovalent XBB.1.5 formulation for the 2023-2024 COVID-19 vaccination campaign.

## Introduction

Prior to recommendations for COVID-19 vaccine strain change and composition harmonization, COVID-19 vaccine options included monovalent SARS-CoV-2 Prototype (Wuhan-Hu-1) and bivalent (Prototype + Omicron BA.4/5). The need to update the vaccine strain has been signaled by a reduction of antibody-mediated immunity against Omicron XBB subvariants [1]. A vaccine strain and composition change to monovalent XBB.1.5-based vaccines have been supported by the World Health Organization (WHO), the European Medicines Agency (EMA), and the United States Food and Drug Administration (FDA) in anticipation of the Fall 2023-2024 booster campaigns [2-4]. This change in vaccine composition highlights the need for robust immunologic evaluation of emerging Omicron XBB sub-variants.

The Novavax COVID vaccine platform contains the full-length recombinant spike (rS) protein presented in its native trimeric and prefusion conformation [5,6]. Spike protein trimers form particles surrounding a polysorbate-80 core, which may improve antigen uptake and processing. Protein antigens are manufactured in a baculovirus expression system using *Spodoptera frugiperda* moth cells that express glycoproteins with truncated N-linked glycans with a potential for enhanced epitope exposure. Antigen particles are formulated with a saponin-based Matrix-M™ adjuvant [7], which has been shown to induce robust antibody responses that protect upper and lower airways in nonhuman primates [8] and polyfunctional Th1-biased CD4+ T cell responses [9-11]. NVX-CoV2373, based on the Prototype (Wuhan-Hu-1) rS, was demonstrated to be well tolerated and immunogenic in humans [11,12], with a vaccine efficacy of 90.4% (95% CI; 82.9 to 94.6) in a Phase 3 clinical trial [13]. NVX-CoV2373 has been authorized for primary series use and as a homologous or heterologous booster in many countries globally.

In 2022, the United States FDA and many global regulators authorized updated bivalent mRNA vaccine boosters containing sequences of both the ancestral and Omicron BA.1 or Omicron BA.4/BA.5 spike proteins. Since the addition of bivalent vaccines, their benefits, compared to monovalent options, have been debated. It may be that the presence of Prototype spike in the current bivalent vaccines leads to original antigen sin, also known as immunological imprinting, that can bias immune responses to the development of lower immunity to the variant spike [14].

Omicron XBB lineages likely emerged following a recombination of two co-circulating Omicron BA.2 lineages, BJ.1 and BM.1.1.1, during the summer of 2022 [15]. At the time of vaccine strain recommendations by the WHO, EMA, and FDA, Omicron XBB.1.5 represented a well characterized strain among the currently prevalent and emergent strains. Sequence comparison supported XBB.1.5 as the preferred vaccine strain due to close sequence homology to other emerging XBB variants. XBB.1.5 has two amino acid changes (E180V and T478R) from XBB.1.16 [16]. XBB.2.3 has three amino acid changes (V252G, D253G, and P521S) compared to XBB.1.5, and five from XBB.1.6 (E180V, G252V, D253G, T478K, P521S). Further, a descendant lineage of XBB.1.9.2, EG.5, has an additional amino acid change (F456L) compared to XBB.1.5. Subvariant EG.5.1, which has an additional amino acid change (Q52H), has become prevalent in some regions of the world [17]. Due to the continued rapid evolution of the SARS-CoV-2 virus, the ability of the updated vaccines to generate cross-protective immunity to future viral variants will be critical as a periodic COVID-19 vaccine strain-change has been suggested. Preclinical data from animal models are needed to inform and support the update of the next-generation COVID-19 vaccine composition.

## Results

### Primary Immunization with Prototype, Bivalent, XBB.1.5, or XBB.1.16 rS in Mice

We assessed humoral immune responses in female BALB/c mice following two-dose primary series immunization with monovalent and bivalent vaccines. Mice (n=10 per group) were inoculated intramuscularly with XBB.1.5 (1 μg rS) or bivalent rS (0.5 μg Prototype rS + 0.5 μg XBB.1.5 rS) on days 0 and 14, and sera were collected at day 21 (1 week after the second dose). The monovalent vaccine appeared to be advantageous compared to a bivalent approach as XBB.1.5 rS elicited comparable cross-reactive pseudovirus neutralizing antibodies against Omicron XBB.1.5 (GMT=4554, 95% CI: 3285-6313), XBB.1.16 (GMT=4156, 95% CI: 2568-6724), and XBB.2.3 (GMT=2058, 95% CI: 1019-4157), and the bivalent vaccine induced a ∼50% or lower response to XBB.1.5 (GMT=981, 95% CI: 625-1541), XBB.1.16 (GMT=2168, 95% CI: 1387-3389), and XBB.2.3 (GMT=524, 95% CI: 224-1227) **(Figure 1A)**.

**Figure 1.**
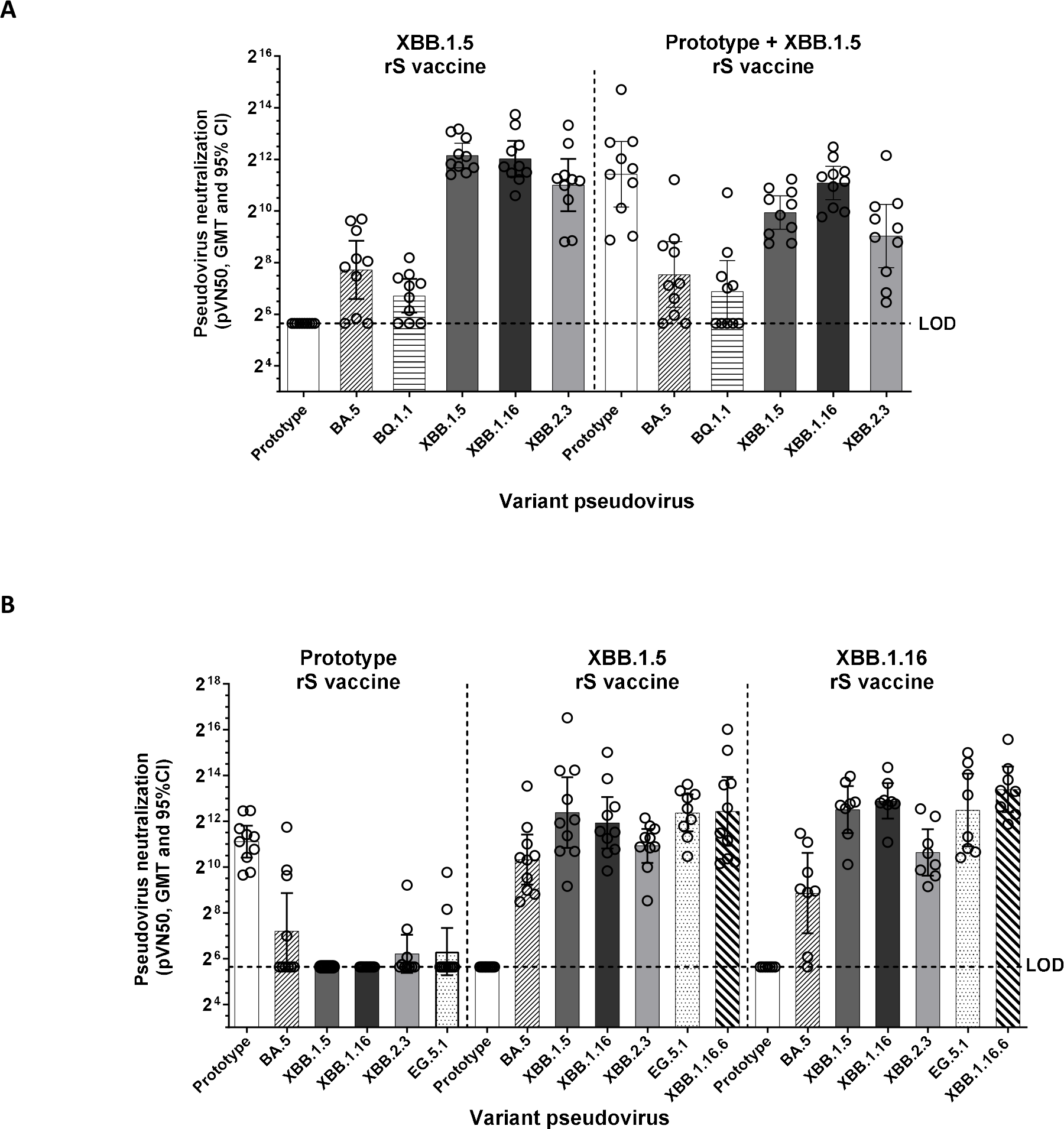
Humoral Responses Following Variant-Adapted Two-Dose Primary Series Vaccination in Mice. (A) Pseudovirus neutralization in mice sera collected one week following primary vaccination with two doses of monovalent XBB.1.5 or bivalent (Prototype + XBB.1.5). (B) Pseudovirus neutralization in mice sera collected one week following primary series vaccination with two doses of monovalent Prototype, XBB.1.5, or XBB.1.16. Open circles represent individual data points, solid bars represent group geometric mean titers, error bars represent 95% confidence intervals, and the horizontal dashed line represents the assay limit of detection (LOD).

We next compared the induction of pseudovirus neutralizing antibodies after immunization with a primary series of various monovalent rS constructs. Groups of mice (n=10) were inoculated intramuscularly with Prototype, XBB.1.5, or XBB.1.16 on days 0 and 14, and sera were collected at day 21 (1 week after the second dose). Humoral responses following XBB.1.5 or XBB.1.16 primary series were comparable and highly immunogenic as they induced pseudovirus neutralizing antibodies against SARS-CoV-2 Omicron BA.5, XBB.1.5, XBB.1.16, XBB.2.3, EG.5.1, and XBB.1.16.6 **(Figure 1B)**. Following the primary vaccination series, homologous variant responses for XBB.1.5 (GMT=4148, 95% CI: 2494-6900) and XBB.1.16 (GMT=7622, 95% CI: 4451-13053) vaccinations were slightly higher than heterologous responses for XBB.1.5 vaccine to XBB.1.16 pseudovirus (GMT=3473, 95% CI: 2282-5287) and XBB.1.16 vaccine to XBB.1.5 pseudovirus (GMT=5846, 95% CI: 2863-11940) **(Figure 1B)**. Compared to homologous variant responses, EG.5.1 neutralization responses were similar in magnitude following immunization with XBB.1.5 (GMT=6478, 95% CI: 3464-12112) and XBB.1.16 (GMT=5770, 95% CI: 1918-17358), and XBB.1.16.6 variant neutralization responses were similar following immunization with XBB.1.5 (GMT=5548, 95% CI: 1960-15704), and XBB.1.16 (GMT=10781, 95% CI: 5413-21472) **(Figure 1B)**. The magnitude of EG.5.1 and XBB subvariant responses were comparable to the levels achieved with homologous responses to a two-dose primary series of Prototype vaccine, suggesting adequate antibody levels were achieved **(Figure 1B)**.

### XBB Lineage Booster Immunization in Mice and Nonhuman Primates

We also investigated immune responses in mice primed with bivalent rS (Prototype rS + BA.5 rS) followed by a booster dose of monovalent XBB.1.5 or XBB.1.16. Groups of mice (n=10 per group) were inoculated intramuscularly with a primary immunization series of a bivalent (Prototype rS + BA.5 rS) vaccine on days 0 and 14, followed by a single booster dose with XBB.1.5 rS or XBB.1.16 rS. Sera were collected at day 21 (1 week after the second dose) and day 61 (2 weeks after booster dose). XBB.1.5 and XBB.1.16 induced a >35-fold increase in pseudovirus neutralizing antibodies against XBB.1.5 and XBB.1.16 compared to the post-priming series day 21 baseline, and were similar in magnitude **(Figure 2A)**. Pseudovirus neutralizing antibodies were higher against XBB.2.3 following immunization with XBB.1.5 (GMT=4561, 95% CI: 2219-9375) compared to levels after immunization with XBB.1.16 (GMT=2938, 95% CI: 1495-5774) **(Figure 2A)**. Pseudovirus neutralizing antibody responses in mice were further analyzed by antigenic cartography. Priming with two doses of bivalent vaccine (Prototype + BA.5) resulted in greater than 30-fold fold-differences between Prototype to both XBB.1.5 and XBB.1.16 **(Figure 2B)**. This large antigenic distance was expected as the neutralized epitopes responses of Prototype and BA.5 rS are largely absent on XBB sub-variants. Antigenic distances with a fold-difference less than 2-fold are considered to be matched responses. Boosting primed mice with XBB.1.5 vaccine induced a matched response to XBB.1.16, with an antigenic distance of 0.691 **(Figure 2B)**. Similarly, an XBB.1.16 boost induced a matched response to XBB.1.5 with a fold-difference of 0.750 **(Figure 2B)**.

**Figure 2.**
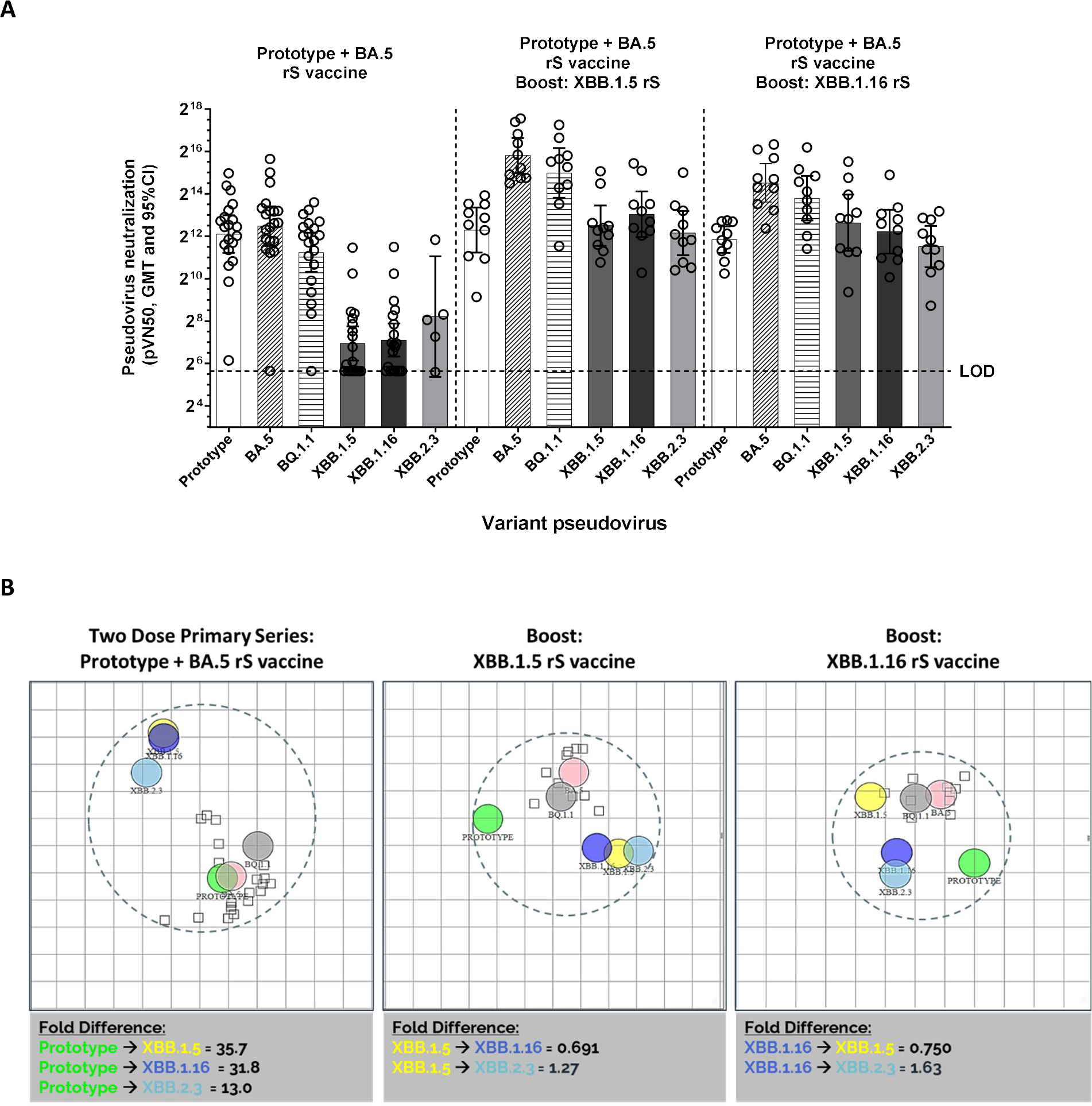
Humoral Responses Following XBB.1.5 Booster in Mice. (A) Pseudovirus neutralization in mice boosted with XBB.1.5 or XBB.1.16 following primary vaccination with bivalent (Prototype + BA.5). Open circles represent individual data points, solid bars represent group geometric mean titers, error bars represent 95% confidence intervals, and the horizontal dashed line represents the assay limit of detection (LOD). (B) Antigenic cartography upon two doses of bivalent Prototype + BA.5 vaccine (left panel, sera collected one week after primary series), following monovalent boost with XBB.1.5 (middle panel), or XBB.1.16 (right panel) vaccines (sera collected two weeks after booster dose). Each square corresponds to one animal and each grid square 1 antigenic distance of 2-fold change in neutralization titer with fold differences given below each map.

The impact of priming series immunization strain and composition on XBB.1.5 pseudovirus neutralization was also tested in nonhuman primates (NHPs). Rhesus macaques (*Macaca mulatta*; n=5 per group) received a two-dose primary series of either Prototype or bivalent (Prototype + BA.5) vaccines and were boosted with XBB.1.5. All NHPs were monitored twice daily to determine the safety of the vaccines, and no local or systemic side effects were reported after the primary series or booster vaccinations. For both priming regimens, boosting with XBB.1.5 induced comparable XBB.1.5, XBB.1.16, XBB.2.3, and EG.5.1 pseudovirus neutralizing responses with overlapping confidence intervals **(Figure 3)**. However, the bivalent primary series regimen containing BA.5 resulted in higher titers against XBB variants compared to titers after a primary series of monovalent Prototype rS **(Figure 3)**.

**Figure 3.**
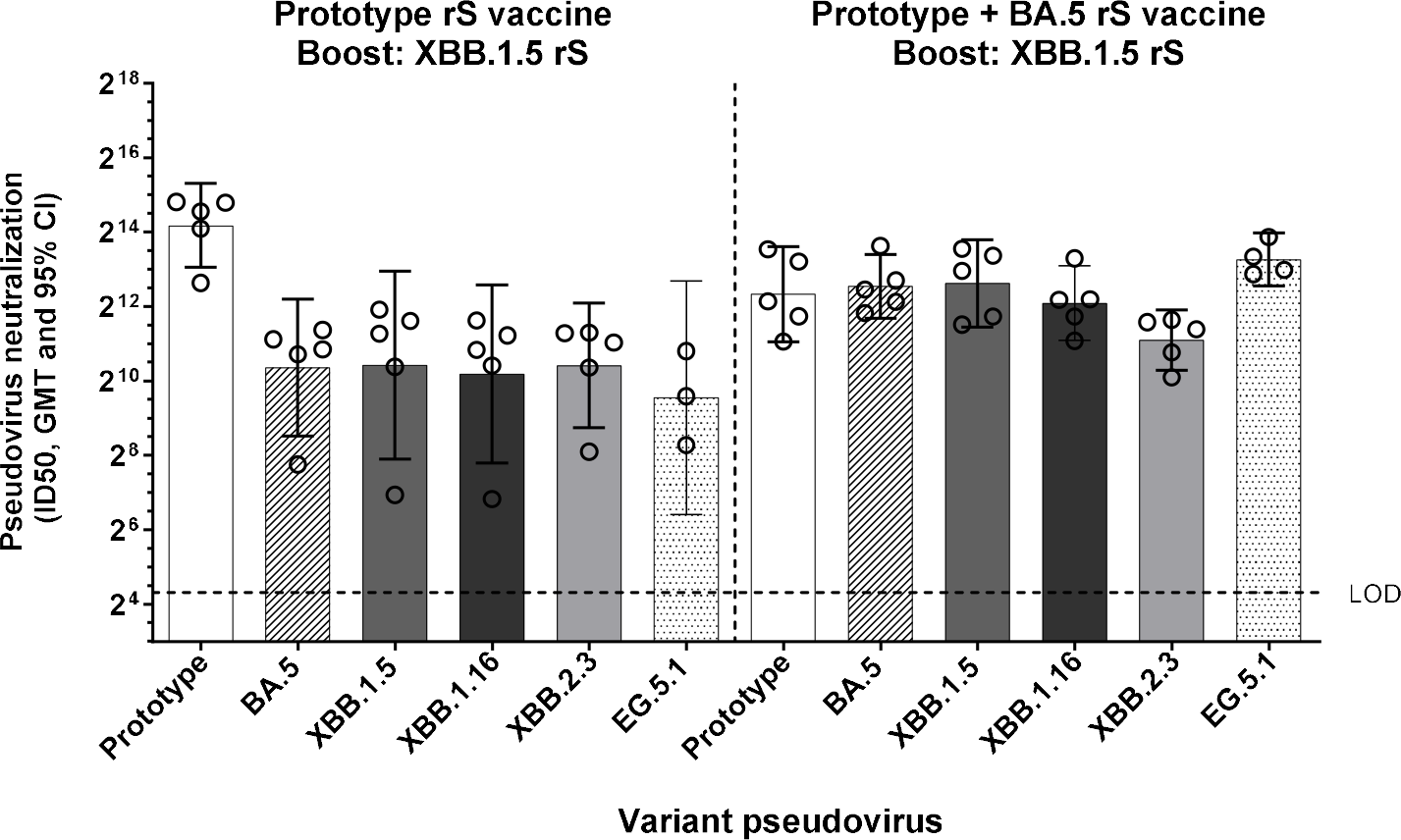
Humoral Responses Following XBB.1.5 Booster in Rhesus Macaques. (A) Pseudovirus neutralization titers in rhesus macaques boosted with XBB.1.5 approximately 8 months after Prototype or bivalent (Prototype + BA.5) priming regimens. Sera were collected two weeks after the booster dose. Open circles represent individual data points, solid bars represent group geometric mean titers, error bars represent 95% confidence intervals, and the horizontal dashed line represents the assay limit of detection (LOD).

### CD4+ T Cell Responses in Mice and Nonhuman Primates

To investigate the cellular responses, we measured CD4+ T cell responses in mice (n=5 per group) immunized with Prototype or bivalent (Prototype + BA.5) priming series vaccine and boosted with XBB.1.5. Th1 (IFN-γ, IL-2, and TNF-α) and Th2 (IL-4) cytokine-producing CD4+ T cells were measured in splenocytes isolated two weeks post booster dose. A robust CD4+ T cell response was recalled at comparable levels post-boost upon stimulation with rS of XBB.1.5, XBB.1.6, or other variants, irrespective of priming vaccine **(Figure 4A)**. Similarly, NHPs primed with bivalent (Prototype + BA.5) vaccine and boosted with XBB.1.5 elicited a Th1-biased cellular response with comparable magnitudes of cytokine-positive cells for all variants tested **(Figure 4B)**.

**Figure 4.**
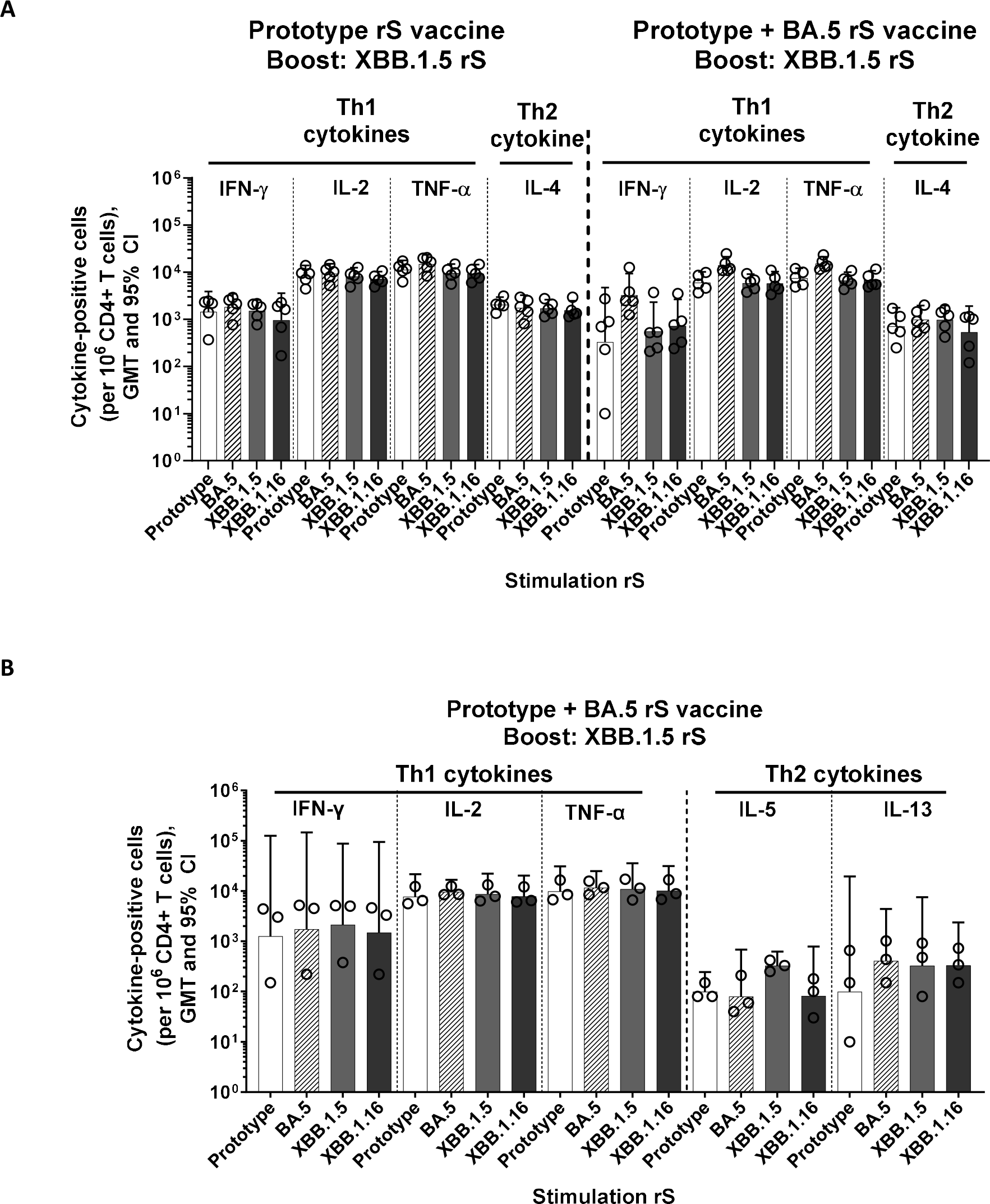
CD4+ T Cell Responses to an XBB.1.5 Booster. (A) CD4+ T cell responses in mice boosted with XBB.1.5 primed with Prototype or bivalent (Prototype + BA.5). Splenocytes were collected two weeks after the booster dose. (B) CD4+ T cell responses in rhesus macaques boosted with XBB.1.5 primed with bivalent (Prototype + BA.5). PBMCs were collected two weeks after the booster dose. Open circles represent individual animal data points, solid bars represent group geometric mean values with 95% CI error bars.

## Discussion

The Novavax monovalent XBB.1.5 vaccine induced cross-neutralizing responses against SARS-CoV-2 Omicron XBB lineage sub-variants including XBB.1.5, XBB.1.16, XBB.2.3, EG.5.1, and XBB.1.16.6. In nonhuman primates, the XBB.1.5 booster was shown to induce similar neutralizing responses against XBB.1.5, XBB.1.16, XBB.2.3, and EG.5.1 in animals primed with monovalent Prototype, or bivalent vaccine. In mice, a primary series of monovalent 1 μg XBB.1.5 rS generated higher-titer neutralizing responses compared to the bivalent vaccine composed of 0.5 μg each of Prototype and BA.5, suggesting the preferential use of a monovalent vaccine. This was likely due to the half a dose of the antigen (0.5 μg of each antigen in the bivalent vaccine compared to 1 μg of antigen in the monovalent vaccine) as there was no indication that original immune imprinting was associated with boosting. The immunogenicity of the monovalent XBB.1.5 vaccine was further evaluated by measuring humoral and cell-mediated immune responses, which showed the presence of functional antibodies that neutralize XBB.1.5, XBB.1.16, XBB.2.3, EG.5.1, and XBB.1.16.6 pseudovirus, and a polyfunctional Th1-biased CD4+ response against XBB sub-variants. These findings are relevant as studies have shown that higher levels of neutralizing antibodies generally correlate with enhanced protection and durability of immune responses [18-24]. The contribution of other mechanisms of immunity is highlighted by the persistence of population-level immunity against severe disease and hospitalization despite reduced levels of vaccine-mediated neutralization of XBB sub-variants. Notably, Th1 cytokine signaling can promote the cytotoxic activities of CD8+ T cells and macrophages to destroy infected cells and limit the severity of disease. Further, Fc-effector functional antibodies induced by NVX-CoV2373 were identified as key determinants of protection against infection in rhesus macaques and humans [25]. Though little focus has been placed on data describing cellular immunity or Fc-effector profiles, their role in a productive adaptive immune response should not be ignored. To our knowledge, this is the first booster study in a non-human primate (NHP) model for evaluating XBB.1.5 booster immunogenicity. This NHP booster immunogenicity study recapitulated the results from the mouse model studies that XBB.1.5 booster produces significant cross-neutralization across the EG.5.1 variant and three major Omicron XBB sub-variants: XBB.1.5, XBB.1.16 and XBB.2.3. Importantly, the XBB.1.5 booster was immunogenic irrespective of priming regimen, as the general population includes individuals primed with diverse vaccination and infection backgrounds. Together, these data demonstrate that a next-generation Novavax COVID-19 vaccine based on monovalent XBB.1.5 rS can induce robust humoral and cellular immunity to EG.5.1 and XBB sub-variants (XBB.1.5, XBB.2.3, and XBB.1.16.6) in mice and NHPs primed with Prototype and bivalent vaccines. Cross-neutralization of the immune response generated by XBB.1.5 across EG.5.1 and other XBB variants (XBB.1.16, XBB.2.3, and XBB.1.16.6) is also encouraging, in addressing emergence of either EG.5.1, XBB.1.16.6, or other strains. Consistent with recommendations by the WHO, EMA, and FDA, our preclinical data in mice and non-human primates support updating the Novavax vaccine to a monovalent XBB.1.5 formulation for the 2023-2024 COVID-19 season.

## Methods

### Vaccine Constructs

SARS-CoV-2 Prototype rS (construct BV2373) was manufactured by the Novavax Discovery Group (Gaithersburg, MD). The SARS-CoV-2 rS vaccine is constructed from the full-length, wild-type SARS-CoV-2 S glycoprotein, based upon the GenBank gene sequence MN90894, nucleotides 21563-25384 (SARS-CoV-2 Wuhan-Hu-1 variant). The native full-length S protein was modified by mutation of the putative furin cleavage site RRAR to QQAQ (3Q) located within the S1/S2 cleavage domain to be protease resistant. Two additional proline amino acid substitutions were inserted at positions K986P and V987P (2P) within the heptad repeat 1 (HR1) domain to stabilize SARS-CoV-2 S in a prefusion conformation, which is believed to optimize presentation of neutralizing epitopes [26].

SARS-CoV-2 Omicron BA.5. XBB.1.5 rS, and XBB.1.16 (constructs BV2540, BV2601, and BV2633), based on the Omicron BA.5, XBB.1.5, and XBB.1.16 variants of SARS-CoV-2, were manufactured by the Novavax Discovery Group (Gaithersburg, MD). Omicron BA.5, XBB.1.5, and XBB.1.16 variant sequences were obtained from the GISAID database references EPI-ISL 12097410.1, 16343574, and 17351426. To produce construct BV2540, the native full-length S protein was subjected to mutations applied to the Prototype Wuhan-Hu-1 rS plus additional mutations: V3G, T19I, A27S, G142D, V213G, G339D, S371F, S373P, S375F, T376A, D405N, R408S, K417N, N440K, L452R, S477N, T478K, E484A, F486V, Q498R, N501Y, Y505H, D614G, H655Y, N679K, P681H, N764K, D796Y, Q954H, and N969K, as well as ΔL24, ΔP25, ΔP26, ΔH69, and ΔV70. To produce construct BV2601, in addition to the mutations applied to the Prototype Wuhan-Hu-1 rS, the following mutations were introduced to the native full-length S protein: T19I, A27S, V83A, G142D, H146Q, Q183E, V213E, G252V, G339H, R346T, L368I, S371F, S373P, S375F, T376A, D405N, R408S, K417N, N440K, V445P, G446S, N460K, S477N, T478K, E484A, F486P, F490S, Q498R, N501Y, Y505H, D614G, H655Y, N679K, P681H, N764K, D796Y, Q954H, and N969K, as well as Δ24-26 and ΔY144. To produce construct BV2633, in addition to the mutations applied to the Prototype Wuhan-Hu-1 rS the following mutations were introduced to the native full-length S protein: K986P, V987P, E180V, K478R from the BV2601 construct. Recombinant baculoviruses were cloned and rS expressed in Sf9 insect cells and purified as described previously [9].

### Animal Ethics Statement

The reporting in this manuscript follows the recommendations in the ARRIVE guidelines.

The mouse studies were conducted at Noble Life Sciences (Sykesville, MD). Animals were maintained and treated according to Animal Welfare Act Regulations, the US Public Health Service Office of Laboratory Animal Welfare Policy on Humane Care and Use of Laboratory Animals, Guide for Care and Use of Laboratory Animals (Institute of Laboratory Animal Resources, Commission on Life Sciences, National Research Council, 1996), and AAALACi accreditation. Mouse studies were approved by Noble Life Sciences IACUC.

The study in rhesus macaques was conducted at Texas Biomedical Research Institute (San Antonio, TX). Animals were maintained at Texas Biomedical Research Institute for the entire in-life portion of the study and were treated according to Animal Welfare Act regulations and the Guide for the Care and Use of Laboratory Animals (2011). Rhesus macaque studies were approved by Texas Biomedical Research Institute IACUC.

### Mouse study designs

For the primary series studies, female BALB/c mice (10-12 weeks old, weight range 17-22 grams, N = 10-20 per group, total 80-100 per study) were immunized by intramuscular (IM) injection with two 1 μg doses spaced 14 days apart (study day 0 and 14) of monovalent Prototype (control), XBB.1.5, XBB.1.16, or bivalent Prototype + XBB.1.5 (0.5 μg each) with 5 μg Matrix-M adjuvant (Novavax, AB, Uppsala, SE). Serum was collected for analysis on study Day 21, one week after the 2^nd^ dose.

For the booster study, female BALB/c mice (N = 10 per group, 200 mice total) were immunized by intramuscular (IM) injection with two 1 μg doses spaced 14 days apart (study day 0 and 14) of Prototype (control) or Prototype + Omicron BA.5 (0.5 μg each) with 5 μg Matrix-M adjuvant. A booster (3^rd^ dose) of 1 μg Omicron XBB.1.5 or XBB.1.16 with 5 μg Matrix-M adjuvant was administered on Day 47 (approximately 1 month post 2^nd^ dose). Sera and spleen were collected 2 weeks after the booster dose on day 61 to evaluate antibody and cellular responses.

For all mouse studies, animals were randomly assigned to groups as they were removed from shipping cages.

### Nonhuman primate study design

Female and male rhesus macaques (*Macaca mulatta*, N = 5 per group; N = 15 total), 3-11 years old and weighing 3-9 kilograms at study initiation, were obtained from a SNPRC specific pathogen free (SPF) colony and/or Envigo (Alice, TX). Animals were randomly assigned to groups based on similar age and sex distribution across the groups. NHPs were immunized by intramuscular injection (0.5 ml) with the human dose level: 5 μg NVX SARS-CoV-2 Prototype (control) or Omicron BA.5 variant rS vaccines adjuvanted with 50 μg Matrix-M administered as monovalent, or bivalent prime/boost on days 0 and 21 (primary series). A booster consisting of Omicron XBB.1.5 rS with 50 μg Matrix-M was administered at week 35 (Day 246). Serum and peripheral blood mononuclear cells (PBMCs) were collected 2 weeks post boost on Day 260.

### Pseudovirus neutralization

SARS-CoV-2 pseudoviruses were generated using a lentivirus platform adapted from Crawford, 2020 [27]. Briefly, backbone and helper plasmids, including Wuhan-Hu-1 spike, were obtained from BEI Resources. Additional variants were synthesized in pcDNA3.1 (GenScript) using the appropriate spike protein sequence from the EPICoV database. All spike protein sequences included a deletion of the cytoplasmic tail. HEK293T cells were seeded one day prior to transfection, incubated at 37°C overnight, and transfected when the cellular monolayer was 60-75% confluent. The transfection uses a cationic-lipid delivery system such as Lipofectamine 3000 (Thermo Fisher) or JetPrime Optimus (Polyplus) with a set of plasmids encoding: a lentiviral backbone, a dual reporter plasmid expressing both luciferase and Zs green, a plasmid expressing SARS-CoV-2 spike (such as Wuhan-Hu-1, Omicron BA.5, and BQ.1.1) and a plasmid expressing other HIV proteins for pseudovirion formation. Then, 48 hours following transfection, supernatants were collected, centrifuged, and filtered through a 0.45 μm filter to obtain a pseudovirus stock. Commercial pseudovirus for Omicron XBB.1.5, XBB.1.16, XBB.2.3, and EG.5.1 were obtained from eEnzyme and incorporated only a luciferase reporter gene for detection of pseudoviral entry. Aliquots of pseudovirus stock were stored at -80°C. All work with pseudovirus was performed in a Biosafety Level 2 laboratory and approved by our Institutional Biosafety Committee.

Each newly produced lot or new shipment of pseudovirus, if commercially obtained, was titered under assay conditions to determine the working dilution to target an RLU of 100,000 prior to testing serum. The pseudovirus neutralization assay was then performed using a HEK293T cell line stably expressing hACE2 (HEK293T/ACE2 obtained from Creative Biogene). Serum samples were heat-inactivated by placing in a 56°C water bath for 30 minutes, followed by cooling to 4°C immediately. Serum samples were serially diluted three-fold in reduced serum Opti-MEM starting at a 1:20 or 1:50 dilution in a 96-well tissue culture plate. Fifty microliters of SARS-CoV-2 Pseudovirus stock (corresponding to 100,000 RLU, range from 50,000-250,000) was then added to each well, followed by incubation at 37°C for one hour. Then, 2.0 × 10^4^ HEK293T/hACE2 cells in 100 μL of HEK293T cell culture medium (DMEM without phenol red + 5% FBS + 1% Penicillin + streptomycin + glutamine) containing 1.25 μg/ml puromycin were added to the wells, followed by incubation for 72 hours at 37°C. After incubation, 50 μL BrightGlo Luciferase Substrate (Promega) was added to each well. Plates were incubated for 5 minutes at room temperature without ambient light. Viral entry into the cells was determined by measuring the luminescence with a SpectraMax iD3 microplate reader. Pseudovirus neutralizing antibody titer of the serum was determined through the absence or reduction of luminescence in a well. Data were analyzed and neutralization curves were generated in GraphPad Prism for each sample; 50% pseudovirus neutralization titers (pVN_50_) and 50% inhibition dilution (ID50) were calculated using 4-parameter curve fitting. No-serum wells were present on each plate along with at least one positive and negative monoclonal antibody for each pseudovirus tested.

A similar pseudovirus neutralization assay, validated for testing human samples (for Ancestral, Omicron BA.5 and XBB.1.5 strains), was utilized as fit-for-purpose for testing NHP samples. This method is similar to the method used for the mouse studies except the input virus targeted an RLU of 50,000 (range 10,000-300,000) per well and diluted in infection medium containing Dulbecco’s Minimal Essential Medium (DMEM) and 2% heat inactivated fetal bovine serum (FBS), test serum and pseudovirus was incubated for 2-hours; 10,000 cells/well were used for the assay. Serum dilution series started at 1:10, which was reported as 1:20 after addition of virus. Luciferase readout was performed from 15 minutes after addition of luciferase reagent up to 60 minutes, followed by data calculation using Softmax 4-parameter curve fit.

### Cellular Assay

For ICCS assay of murine splenocytes, cells were cultured in a 96-well U-bottom plate at 1-2 × 10^6^ cells per well. The cells were stimulated with NVX-CoV2373 or the indicated SARS-CoV-2 variant spike protein.

The plate was incubated 6 hours at 37 °C in the presence of BD GolgiPlug™ and BD GolgiStop™ (BD Biosciences) for the last 4 hours of culture. Cells were labeled with murine antibodies BV650 CD3 (Clone 145-2C11, 1:25), APC-H7 CD4 (Clone GK1.5, 1:25), FITC CD8 (Clone 53-6.7, 1:25), Alexa Fluor 700 CD44 (Clone IM7, 1:50), and PE CD62L (Clone MEL-14, 1:50) (BD Pharmingen, CA), and the yellow LIVE/DEAD® dye (1:300). After fixation with Cytofix/Cytoperm (BD Biosciences), cells were incubated with PerCP-Cy5.5-conjugated anti-IFN-γ (Clone XMG1.2, 1:50), BV421-conjugated anti-IL-2 (Clone JES6-5H4, 1:100), PE-cy7-conjugated anti-TNF-α (Clone MP6-XT22, 1:800), and APC-conjugated anti-IL-4 (Clone 11B11, 1:100) (BD Biosciences). All stained samples were acquired using an LSR-Fortessa flow cytometer or Symphony A3 (Becton Dickinson, San Jose, CA) and the data were analyzed with FlowJo software version 10 (Tree Star Inc., Ashland, OR). Data shown were gated on CD44^hi^ CD62L^low^ effector CD4+ T cell population.

For ICCS assay of NHP PBMCs, the cells were thawed and rested at 37 °C overnight. The cells were then stimulated as described above with NVX-CoV2373 or the indicated variant protein. Cells were labeled with human/NHP antibodies BV650-conjugated anti-CD3 (Clone SP34-2, 1:10), APC-H7-conjugated anti-CD4 (Clone L200, 1:10), APC-conjugated anti-CD8 (Clone RPA-T8, 1:10), and the yellow LIVE/DEAD® dye (1:300) for surface staining; BV421-conjugated anti-IL-2 (Clone MQ1-17H12, 1:25), PerCP-Cy5.5-conjugated anti-IFN-γ (Clone 4S. B3, 1:10), and PE-cy7-conjugated anti-TNF-α (Clone Mab11, 1:50) (BD Biosciences) for intracellular staining. Data shown were gated on CD4+ T cell population.

### Statistical Analysis

Geometric mean titer (GMT) and 95% confidence interval (95% CI) were calculated with GraphPad Prism 9.0 software (La Jolla, CA) from serum antibody titers and plotted.

## Additional Information

Registered and proprietary names, e.g., trademarks, used herein are not to be considered unprotected by law even when not specifically marked as such.

## Acknowledgements

Medical writing and editing support were provided by Ashfield MedComms (New York, USA), an Inizio company, and funded by Novavax, Inc.

## Author Contributions

Conceptualization: N.P., R.M.M., F.D., G.S. Investigation: N.P., J.F.T., M.G., H.Z., J.N., D.J., Z.C., M.Z., R.K., G.S. Data Analysis: N.P., R.K. Writing—original draft: A.M.M. Writing—review and editing: N.P., J.F.T., M.G., H.Z., A.M.M., A.M.G., R.M.M., R.K., F.D., G.S. Visualization: A.M.G., N.P. Project Administration: M.G.

## Data Availability

The datasets generated during and/or analyzed during the current study are available from the corresponding author on reasonable request.

